# Ataxia Telangiectasia triggers deficits in Reelin pathway

**DOI:** 10.1101/336842

**Authors:** Júlia Canet-Pons, Ralf Schubert, Ruth Pia Duecker, Roland Schrewe, Sandra Wölke, Martina Schnölzer, Georg Auburger, Stefan Zielen, Uwe Warnken

## Abstract

Autosomal recessive Ataxia Telangiectasia (A-T) is characterized by radiosensitivity, immunodeficiency and cerebellar neurodegeneration. A-T is caused by inactivating mutations in the Ataxia-Telangiectasia-Mutated (ATM) gene, a serine-threonine protein kinase involved in DNA-damage response and excitatory neurotransmission. The selective vulnerability of cerebellar Purkinje neurons (PN) to A-T is not well understood.

Employing global proteomic profiling of cerebrospinal fluid from patients at ages around 15 years we detected reduced Calbindin, Reelin, Cerebellin-1, Cerebellin-3, Protocadherin Fat 2, Sempahorin 7A and increased Apolipoprotein -B, -H, -J peptides. Bioinformatic enrichment was observed for pathways of chemical response, locomotion, calcium binding and complement immunity. This seemed important, since secretion of Reelin from glutamatergic afferent axons is crucial for PN radial migration and spine homeostasis. Reelin expression is downregulated by irradiation and its deficiency is a known cause of ataxia. Validation efforts in 2-month-old *Atm*-/- mice before onset of motor deficits confirmed transcript reductions for Reelin receptors *Apoer2/Vldlr* with increases for their ligands *Apoe/Apoh* and cholesterol 24-hydroxylase *Cyp46a1*. Concomitant dysregulations were found for *Vglut2/Sema7a* as climbing fiber markers, glutamate receptors like *Grin2b* and calcium homeostasis factors (*Atp2b2, Calb1, Itpr1*), while factors involved in DNA damage, oxidative stress, neuroinflammation and cell adhesion were normal at this stage.

These findings show that deficient levels of Reelin signaling factors reflect the neurodegeneration in A-T in a sensitive and specific way. As an extracellular factor, Reelin may be accessible for neuroprotective interventions.

## Introduction

Ataxia-telangiectasia (A-T) is an autosomal recessive disorder defined clinico-pathologically by degeneration of cerebellar Purkinje neurons (PN) combined with a dilatation of oculocutaneous small blood vessels. The patients suffer from immunodeficiency and radiosensitivity, which predisposes them to cancer [1]. A specific diagnostic biomarker in the extracellular compartment, the elevation of alpha-fetoprotein in blood serum is usually detected early in the disease progression [2,3]. The pathology is caused by mutations in the *ATM* (Ataxia-Telangiectasia Mutated) protein [4], a serine-threonine kinase that mainly participates in the DNA double-strand break response [5-8].

Unfortunately, none of its phosphorylation targets are among the critical factors of cerebellar ataxia pathways, so it remains to be understood how the vulnerability to DNA damage connects with the preferential neurodegeneration of Purkinje cells. Oxidative stress has been proposed as crucial, in view of several observations in patients and mouse models [9]. In A-T patient blood, elevated oxidative damage to lipids and DNA as well as altered glutathione levels were observed, suggesting adaptive mechanisms to oxidative stress [10,11]. A-T blood lymphocytes and skin fibroblasts show enhanced vulnerability to oxidative stress [12,13]. However, exposure of patient blood cell cultures to ionizing radiation or bleomycin showed significant differences in chromosomal breaks, while no elevation of oxidative stress markers TBARS, CAT, SOD was detected [14]. A re-analysis of this question in model organisms upon the first description of *Atm*-/- mice has emphasized that affected organs are targets of ROS-mediated damage, in particular the cerebellar PN [15]. The prominent role of oxidative stress for the deficient survival and dendritogenesis of PN was independently confirmed in *Atm*-/- mice [16]. Accumulation of DNA damage, alterations of the redox status and deficient calcium-dependent excitability in the brains of *Atm*-/- mice could be demonstrated [17-19]. However, these investigations are hampered by the early death of *Atm*-/- mice by the age of 4-6 months due to thymic lymphomas, at a stage when their cerebellum appears histologically normal and the mice have no relevant movement deficit [20]. Only when their survival is extended to ages around 8 months by bone-marrow transplantation, a significant cerebellar atrophy with reduced numbers of PN becomes apparent by magnetic resonance imaging and calbindin-immunohistochemistry [21].

To identify further molecular biomarkers of risk and progression in A-T in the extracellular fluid, we studied cerebrospinal fluid (CSF) from patients by global proteome profiling with label-free mass-spectrometry (MS). Several pathway dysregulations were observed, which reflect deficient Reelin signals as completely novel finding, and altered calcium homeostasis, cellular motility and adhesion, as well as neuroinflammation. The crucial effects were validated at transcript level in the *Atm*-/- cerebellar tissue at very young age to assess their robustness and sensitivity. Importantly, we confirm very early and prominent deficits in Reelin signaling, which is secreted from cerebellar glutamatergic afferent fibers to control PN radial migration and dendritic differentiation.

## Materials and methods

### Patients

Cerebrospinal fluid (CSF) was collected from 12 patients with A-T (7 male and 5 female) with an age distribution from 2.6 to 16.2 years (mean 10 years) and 12 control individuals (8 male and 4 female) that were age matched (0.2 to 19.6, mean 8.6 years) (Table 1). The A-T patients were diagnosed on the basis of clinical criteria and AFP values, according to recent World Health Organization (WHO) recommendations [22]. The control group consisted of non-A-T patients receiving lumbar puncture due to another clinical indication. The SARA score [23] was used in patients as a quantitative measurement of the cerebellar ataxia.

**Table 1.**
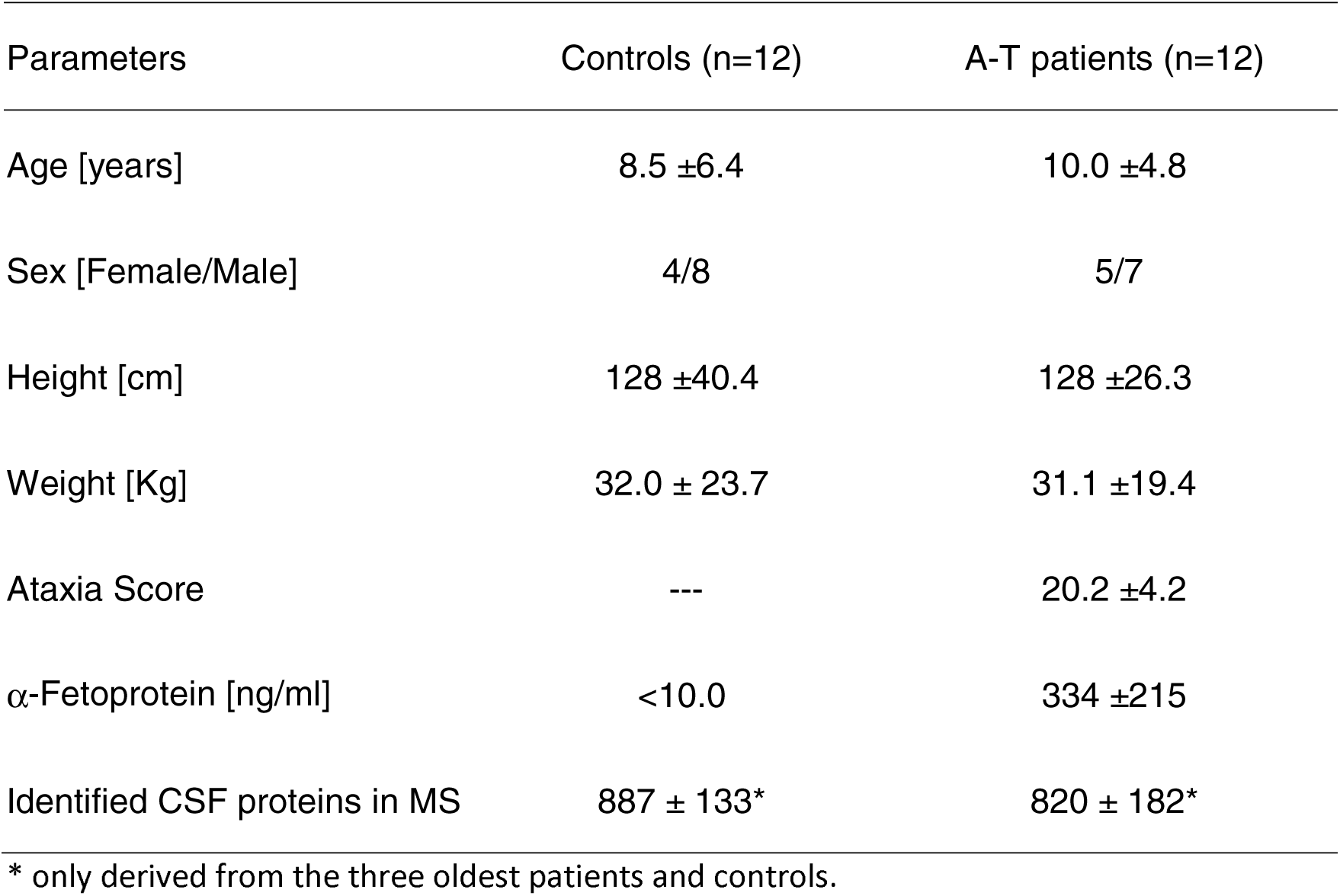
Patients’ Characteristics Basal diagnostic features of individuals studied.

| Parameters | Controls (n=12) | A-T patients (n=12) |
| --- | --- | --- |
| Age [years] | 8.5 ± 6.4 | 10.0 ± 4.8 |
| Sex [Female/Male] | 4/8 | 5/7 |
| Height [cm] | 128 ± 40.4 | 128 ± 26.3 |
| Weight [Kg] | 32.0 ± 23.7 | 31.1 ± 19.4 |
| Ataxia Score | --- | 20.2 ± 4.2 |
| α-Fetoprotein [ng/ml] | <10.0 | 334 ± 215 |
| Identified CSF proteins in MS | 887 ± 133* | 820 ± 182* |

\* only derived from the three oldest patients and controls.

Written consent from patients or caregivers was obtained from each subject. The study was conducted following the ethical principles of the Declaration of Helsinki, regulatory requirements and the code of Good Clinical Practice. The study was approved by the responsible ethics committees (application number 296/09) in Frankfurt and registered at clinicaltrials.gov NCT02285348.

Proteomic analysis of the CSF derived from the three oldest patients (14 years, 3 months; 14 years, 11 months and 16 years, 2 months; 2 male and 1 female) and the three oldest controls (14 years; 15 years, 7 months and 16 years, 9 months; 2 male and 1 female) were selected for further CSF analyses, which was performed by a label-free protein quantification approach using nano ultra-high performance liquid chromatography/ nano electrospray mass spectrometry (nano UPLC/ nanoESI-MS). Analysis of the data was performed with the MaxQuant quantitative proteomics software.

### Sample preparation, gel electrophoresis and tryptic digestion

Protein amounts ranging from 17 to 27 µg (see below) were precipitated according to established protocols [24]. The obtained pellets were incubated with 30 µl of LDS-buffer containing reducing agent (Invitrogen, Carlsbad, USA) for 10 min at 70 °C. Subsequently, proteins were separated by 1D gel electrophoresis on a NuPAGE 4-12% Bis-Tris gradient gel (Invitrogen, Carlsbad, USA) using a MOPS-buffer system. The gel was stained with colloidal Coomassie Blue for 3 h. Afterwards each lane was cut into 27 pieces. Each individual gel piece was washed once with 100 µl H_2_O and 100 µl H_2_O/acetonitrile 50/50 (v/v) and incubated for 5 min at 37 °C. After removing the solution from the gel plugs proteins were alkylated with 100 µl 55 mM iodoacetamide in 40 mM NH_4_HCO_3_ for 30 min at 25 °C in the dark, followed by three alternating washing steps each with 150 µl of water and H_2_O/acetonitrile 50/50 (v/v) for 10 min at 37 °C. Gel pieces were dehydrated with 100 µl neat acetonitrile for 1 min at room temperature, dried for 15 min and subsequently rehydrated with porcine trypsin (sequencing grade, Promega, Mannheim, Germany) with a minimal volume sufficient to cover the gel pieces after rehydration (100 ng trypsin in 40 mM NH_4_HCO_3_). After overnight digestion the supernatant was collected in PCR tubes, while gel pieces were subjected to four further extraction steps (acetonitrile/0.1% TFA 50/50 (v/v)). The combined solutions were evaporated to dryness in a speed-vac concentrator and redissolved in 0.1% TFA/2.5% hexafluoroisopropanol and subsequently analyzed by nanoLC-ESI-MS/MS.

### NanoLC ESI-MS/MS analysis

Tryptic peptides mixtures were separated using a nano Acquity UPLC system (Waters GmbH, Eschborn, Germany). Peptides were trapped on a nano Acquity C18 column, 180 µm x 20 mm, particle size 5 µm (Waters GmbH, Eschborn, Germany). The liquid chromatography separation was performed on a C18 column (BEH 130 C18 100 µm x 100 mm, particle size 1.7 µm (Waters GmbH, Eschborn, Germany) with a flow rate of 400 nl /min. Chromatography was carried out using a 1h gradient of solvent A (98.9% water, 1% acetonitrile, 0.1% formic acid) and solvent B (99.9% acetonitrile and 0.1% µl formic acid) in the following sequence: from 0 to 4% B in 1 min, from 4 to 40% B in 40 min, from 40 to 60% B in 5 min, from 60 to 85% B in 0.1 min, 6 min at 85% B, from 85 to 0% B in 0.1 min, and 9 min at 0% B. The nanoUPLC system was coupled online to an LTQ Orbitrap XL mass spectrometer (Thermo Scientific, Bremen, Germany). The mass spectrometer was operated in the data-dependent mode to automatically measure MS1 and MS2. Following parameters were set: ESI voltage 2400 V; capillary temperature 200 °C, normalized collision energy 35 V. Data were acquired by scan cycles of one FTMS scan with a resolution of 60000 at m/z 400 and a range from 300 to 2000 m/z in parallel with six MS/MS scans in the ion trap of the most abundant precursor ions.

### MaxQuant

Protein quantification was performed with the MaxQuant software 1.5.3.8 [25] wherein peptide identification was performed using the Andromeda [26] search engine integrated into the MaxQuant environment against the human SwissProt database (uniprot-organism—9606+reviewed—yes 03/2016, 20274 sequences). The peptide mass tolerance for database searches was set to 7 ppm and fragment mass tolerance to 0.4 Da. Cysteine carbamidomethylation was set as fixed modification. Variable modifications included oxidation of methionine, deamidation of asparagine and glutamine, as well as protein N-terminal acetylation. Two missed cleavage sites in case of incomplete trypsin hydrolysis were allowed. Furthermore, proteins were considered as identified if at least two unique peptides were identified. Identification under the applied search parameters refers to false discovery rate (FDR) <1% and a match probability of p<0.01, where p is the probability that the observed match is a random event. Data transformation and evaluation was performed with the Perseus software (version 1.5.2.4) [27]. Contaminants as well as proteins identified by site modification only and proteins derived from decoy database containing reversed protein sequences were strictly excluded from further analysis. Protein ratios were calculated by label free quantification (LFQ) comparing experiment and control samples. Filtering for quantitative values was applied to have at least two valid LFQ values in three replicates of either the experiment or the control group. To avoid zero LFQ values for calculating protein expression levels, values equal zero were substituted by values calculated from normal distribution of each data set regarded. For statistical analyses, 2-sample t-tests were performed to calculate differences in the protein abundance between these groups (p-value <0.05). Proteins were only considered as significantly regulated and ranked as potential PPI complex partners if their abundance changed more than +2-fold.

### Bioinformatic pathway enrichment analyses

For protein-protein interaction (PPI) network analysis, the software tool String v.10 (https://string-db.org/) with standard settings has been employed to visualize networks of significant dysregulations [28]. As recommended, gene symbols of factors with significant dysregulation were entered into the Multiple Proteins window, so that the graphic interaction diagram was generated and archived. Automated network statistics were done; significant functional enrichments of GO (Gene Ontology) terms and KEGG pathways were exported into EXCEL files.

### Animal breeding and brain dissection

*Atm*-/- (*Atm*^tm1Awb^) mice were used as the model of A-T [15], together with age- and sex-matched wildtype (WT) animals in the 129SvEv genetic background strain. All procedures were performed according to protocols approved by the German Animal Subjects Committee (Gen.Nr.FK/1006). Mice were housed in accordance with the German Animal Welfare Act, Council Directive of 24 November 1986 (86/609/EWG) Annex II, ETS123, and the EU Directive 2010/63/EU for animal experiments at the FELASA-certified Central Animal Facility (ZFE) of the Frankfurt University Medical School. The mice were housed in Type II L cages (365×207×140 mm^3^, floor area 530 cm^2^), with mutants and controls being bred and aged in parallel under controlled conditions of temperature, humidity, and light/ dark cycles of 12 h. Food and water were accessible *ad libitum*. Genotyping was performed on extracted tail DNA using PCR techniques were described previously [29]. Following cervical dislocation, fresh cerebellar tissue from 8 *Atm*-/- versus 8 WT animals with an age of 1-3 months was dissected and snap-frozen in liquid nitrogen for expression analyses.

### Expression study at transcript level in a mouse model of A-T

Total RNA from frozen cerebellum was isolated using TRIzol reagent (Sigma-Aldrich), following the instructions of the manufacturer. The RNA quality and quantity were measured with a BioPhotometer (Eppendorf) at 260 nm. DNase treatment of 1 µg RNA was performed using DNase Amplification Grade (Invitrogen). Reverse transcription was carried out with SuperScript IV VILO Master Mix (Invitrogen) as recommended. Gene expression assays were performed from 20 ng of cDNA in 96-well optical plates with a StepOnePlus Real-Time PCR system. Replicates from each sample were included per assay. The reaction mix consisted of 5 µl cDNA, 1 µl 20x TaqMan gene expression assay, 10 µl of 2x FastStart Universal Probe Master (ROX; Roche) and 4 µl Gibson DNAse/RNAse free water. The following TaqMan assays were used for reverse-transcriptase quantitative polymerase chain reaction (qPCR) in the mouse tissue: *Abca1* - Mm00442646_m1; *Apob* - Mm01545150_m1; *Apod* - Mm01342307_m1; *Apoe* - Mm00437573_m1; *Apoer2* - Mm00474030_m1; *Apoh* - Mm00496516_m1; *Apoj* (*Clu*) - Mm00442773_m1; *Atp2a2* - Mm01201431_m1; *Atp2b2* - Mm00437640_m1; *Atr* - Mm01223626_m1; *Calb1* - Mm004886647_m1; *Calm1* - Mm01336281_g1; *Cbln1* - Mm01247194_g1; *Cbln3* - Mm00490772_g1; *Cntn2* - Mm00516138_m1; *Cyp46a1* - Mm00487306_m1; *Dab1* - Mm00438366_m1; *Dab2* - Mm01307290_m1; *Efemp1* - Mm01295779_m1; *Fan1* - Mm00625959_m1; *Fat2* - Mm01295779_m1; *Fgfr1* - Mm00438930_m1; *Gria1* - Mm00433753_m1; *Gria2* - Mm00442822_m1; *Gria3* - Mm00497506_m1; *Gria4* - Mm00444754_m1; *Grin1* - Mm00433800_m1; *Grin2a* - Mm00433802_m1; *Grin2b* - Mm00433820_m1; *Grin2c* - Mm00439180_m1; *Grin2d* - Mm00433822_m1; *Grin3a* - Mm01341722_m1; *Grin3b* - Mm00504568_m1; *Grm1* - Mm00810219_m1; *Grm3* - Mm00725298_m1; *Grm4* - Mm01306128_m1; *Grm5* - Mm00690332_m1; *Grm7* - Mm01189424_m1; *Grm8* - Mm00433840_m1; *Gsk3b* - Mm00444911_m1; *Gsr* - Mm00439154_m1; *Gss* - Mm00515065_m1; *Gstp1* - Mm04213618_gH; *Hmgcs1* - Mm01304569_m1; *Homer3* - Mm00803747_m1; *Ighm* - Mm01718955_g1; *Inpp5a* - Mm00805812_m1; *Itih2* - Mm01337594_m1; *Itih4* - Mm00497648_m1; *Itpka* - Mm00525139_m1; *Itpr1* - Mm00439907_m1; *Jup* - Mm00550256_m1; *KitL* - Mm00442972_m1; *Ldlr* - Mm01177349_m1; *Lrp1* - Mm00464608_m1; *Lrp6* - Mm00999795_m1; *Mlh1* - Mm00503449_m1; *Pafah1b1* - Mm00443070_m1; *Pafah1b2* - Mm00476594_m1; *Pafah1b3* - Mm00476597_m1; *Pms2* - Mm01200871_m1; *Reln* - Mm00465200_m1; *Rrm2b* - Mm01165706_m1; *Scg2* - Mm04207690_m1; *Sema7a* - Mm00441361_m1; *Sod1* - Mm01344233_g1; *Sparc* - Mm00486332_m1; *Trpc3* - Mm00444690_m1; *Tst* - Mm01195231_m1; *Txnb* - Mm00466624_m1; *Vglut1* - Mm00812886_m1; *Vglut2* (*Slc17a6*) - Mm00499876- m1; *Vldlr* - Mm00443298_m1. Transcript levels of *Tbp* (Mm00446973_m1) were employed as internal loading control for normalization. Expression levels were analyzed with the 2^−ΔΔCt^ method [30].

## Results and Discussion

### A-T patient CSF shows progressive elevation of protein and albumin

To identify diagnostic markers and to elucidate the molecular events for the neurodegeneration in A-T, we analyzed the CSF from 12 patients and 12 sex- and age-matched control individuals. In addition to diagnostic tests such as neuroimaging(Fig. 1A), clinical ataxia scores and determination of alpha-fetoprotein levels were documented (summary of basal characteristics in Table 1). In good correlation with the progressive increase of ataxia scores during the ageing process (Fig. 1B), also the CSF total protein and albumin levels increased significantly in patients, but not in controls (Fig. 1C-D). Among patients aged 10 to 16 years, total protein and albumin levels showed better correlation to age than the ataxia scores.

**Figure 1.**
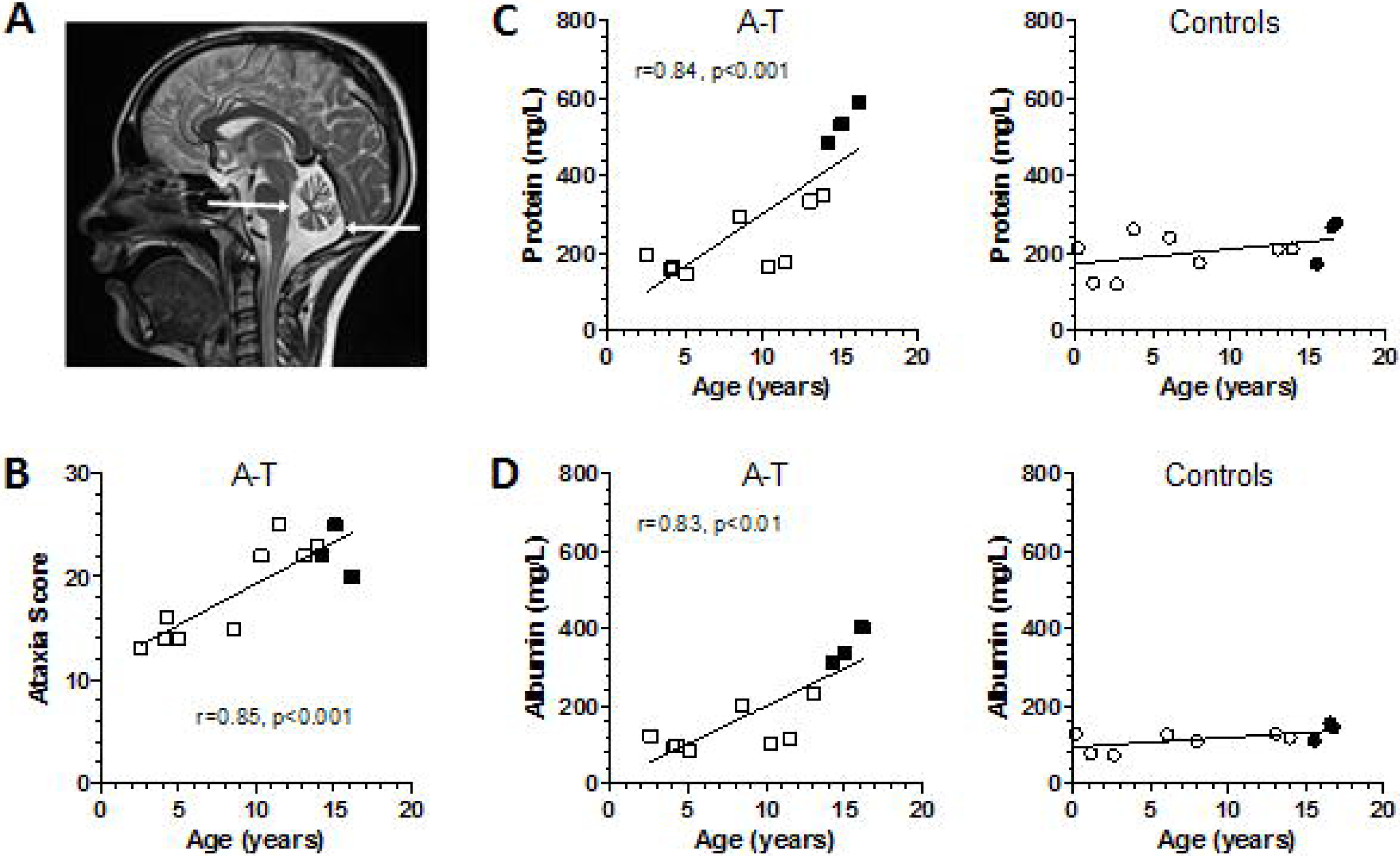
Progressive neurodegeneration in A-T: **(A**) The shrinking cerebellum of an A-T patient is surrounded by excess amounts of CSF (highlighted by white arrows). This extracellular fluid generates high intensity values (shown as light grey or white) in this T2-weighted magnetic resonance image. **(B)** Age-associated progression of SARA ataxia scores is documented for 12 patients. **(C)** Total protein and **(D)** Albumin concentrations analyzed in CSF of the 12 A-T patients and control individuals. Black symbols illustrate the 3 patients and 3 controls selected for CSF global proteome profiling.

### Global proteomics reveals several dysregulations in the Reelin signaling pathway

The three oldest patients (ages 14-16 years) with strongest increase of total protein and albumin were selected for further CSF analyses by MS, together with the three oldest controls. The global proteome bioinformatics identified similar numbers of CSF proteins in patient and controls (Table 1). In comparison to previously reported data [31], we identified 820 proteins in the CSF samples of our group of patients, among which 66 proteins were found with significant differences in protein expression, including downregulations for 44 proteins and upregulations for 22 proteins (Table 2). Our data confirm all the 13 priority proteins reported previously in the CSF of A-T patients [31], but only Secretogranin and SPARC dysregulations reached significance in CSF of our patients.

**Table 2.** Proteins differentially expressed in CSF in the three oldest patients with A-T vs. control individuals.

| Protein Names | Gene Names | Minus LOG<br>(P-value) | Fold Change<br>(negative values<br>indicate down<br>regulations) |
| --- | --- | --- | --- |
| Carbonic anhydrase 2 | CA2 | 1.46 | 6.03 |
| Ig mu chain C region | IGHM | 1.44 | 4.47 |
| C4b-binding protein alpha chain | C4BPA | 2.93 | 3.84 |
| Adenylate kinase isoenzyme 1 | AK1 | 2.17 | 3.73 |
| Apolipoprotein B-100;Apolipoprotein B-48 | APOB | 2.36 | 3.68 |
| Carboxypeptidase N subunit 2 | CPN2 | 1.46 | 3.03 |
| CD5 antigen-like | CD5L | 2.19 | 2.95 |
| Eukaryotic translation initiation<br>Factor 5A and Like1 | EIF5A; EIF5AL1 | 1.60 | 2.66 |
| Chitinase-3-like protein 2 | CHI3L2 | 1.43 | 1.94 |
| Isocitrate dehydrogenase [NADP]<br>cytoplasmic | IDH1 | 1.68 | 1.69 |
| Inter-alpha-trypsin inhibitor heavy<br>chain H1 | ITIH1 | 1.92 | 1.44 |
| Neurogenic locus notch homolog<br>protein 3 | NOTCH3 | 1.46 | 1.38 |
| Serum amyloid P-component | APCS | 3.47 | 1.31 |
| Inter-alpha-trypsin inhibitor heavy chain H2 | ITIH2 | 1.78 | 1.24 |
| Serum paraoxonase/arylesterase 1 | PON1 | 1.41 | 1.19 |
| Junction plakoglobin | JUP | 2.01 | 1.12 |
| Inter-alpha-trypsin inhibitor heavy chain H4 | ITIH4 | 2.39 | 1.09 |
| Complement component C6 | C6 | 1.71 | 0.99 |
| Complement component C8 gamma chain | C8G | 1.30 | 0.96 |
| Clusterin | CLU | 1.60 | 0.67 |
| Beta-2-glycoprotein 1 | APOH | 1.40 | 0.65 |
| 14-3-3 protein zeta/delta | YWHAZ | 1.31 | 0.59 |
| Contactin-2 | CNTN2 | 1.74 | -0.36 |
| Fibulin-1 | FBLN1 | 2.19 | -0.41 |
| Gamma-glutamyl hydrolase | GGH | 2.00 | -0.66 |
| SPARC-like protein 1 | SPARCL1 | 1.60 | -0.66 |
| Plexin-B2 | PLXNB2 | 1.35 | -0.69 |
| 45kDa calcium-binding protein | SDF4 | 2.31 | -0.70 |
| Beta-mannosidase | MANBA | 1.45 | -0.79 |
| Pro-cathepsin H | CTSH | 1.63 | -0.83 |
| EGF-containing fibulin-like extracellular<br>matrix protein 1 | EFEMP1 | 2.44 | -1.02 |
| Secreted Protein Acidic And Cysteine Rich | SPARC | 3.02 | -1.04 |
| Ectonucleotide pyrophosphatase/<br>phosphodiesterase family member 2 | ENPP2 | 1.32 | -1.06 |
| N-acetylglucosamine-1-phosphotransferase<br>subunit gamma | GNPTG | 1.50 | -1.06 |
| 78 kDa glucose-regulated protein | HSPA5 | 1.67 | -1.06 |
| Pyruvate kinase PKM | PKM; PKM2 | 1.47 | -1.15 |
| Di-N-acetylchitobiase | CTBS | 1.85 | -1.19 |
| Collagen alpha-1(VI) chain | COL6A1 | 2.87 | -1.20 |
| Lymphocyte antigen 6H | LY6H | 1.42 | -1.20 |
| Dihydropteridine reductase | QDPR | 1.66 | -1.32 |
| Semaphorin-7A | SEMA7A | 1.31 | -1.34 |
| Secretogranin-1 | CHGB | 2.69 | -1.34 |
| Fibroblast growth factor receptor 1 | FGFR1 | 2.21 | -1.38 |
| Cell surface glycoprotein MUC18 | MCAM | 1.68 | -1.39 |
| Glia-derived nexin | SERPINE2 | 1.58 | -1.57 |
| Carbonic anhydrase 4 | CA4 | 1.50 | -1.71 |
| Phosphoinositide-3-kinase-interacting<br>protein 1 | PIK3IP1 | 1.37 | -1.86 |
| Probable serine carboxypeptidase CPVL | CPVL | 1.43 | -1.89 |
| Carboxypeptidase Q | CPQ | 2.80 | -2.03 |
| Phosphatidylcholine-sterol acyltransferase | LCAT | 1.37 | -2.33 |
| Mammalian ependymin-related protein 1 | UCC1; EPDR1 | 2.17 | -2.37 |
| Fibromodulin | FMOD | 2.16 | -2.38 |
| Cerebellin-1 | CBLN1 | 2.25 | -2.42 |
| Kallikrein-11 | KLK11 | 2.68 | -2.42 |
| Mannosyl-oligosaccharide 1.2-alpha-<br>mannosidase IA | MAN1A1 | 2.41 | -2.43 |
| Reelin | RELN | 2.21 | -2.69 |
| Contactin-6 | CNTN6 | 1.58 | -2.71 |
| Plasma alpha-L-Fucosidase | FUCA2 | 2.33 | -2.75 |
| Calbindin | CALB1 | 2.24 | -2.75 |
| Collectin-12 | COLEC12 | 1.58 | -2.98 |
| Nucleobindin-1 | NUCB1 | 2.69 | -3.13 |
| Testican-2 | SPOCK2 | 1.99 | -3.85 |
| Complement C1q tumor necrosis factor-<br>related protein 5 | C1QTNF5 | 4.40 | -3.85 |
| Cerebellin-3 | CBLN3 | 1.99 | -4.65 |
| Protocadherin Fat 2 | FAT2 | 1.47 | -5.09 |
| Ig gamma-4 chain C region | IGHG4 | 1.51 | -5.14 |

The visualization of our data in volcano plots (Fig. 2), where fold changes are plotted on the X-axis and significance is plotted on the Y-axis, revealed the prominent PN vulnerability by a particularly strong reduction of Calbindin (Log_2_ fold change of −2.75 representing a decrease to 15%), as expected for this abundant calcium-homeostasis factor that is specific for the GABA-ergic PN and is usually deficient in cerebellar ataxias [32].

**Figure 2.**
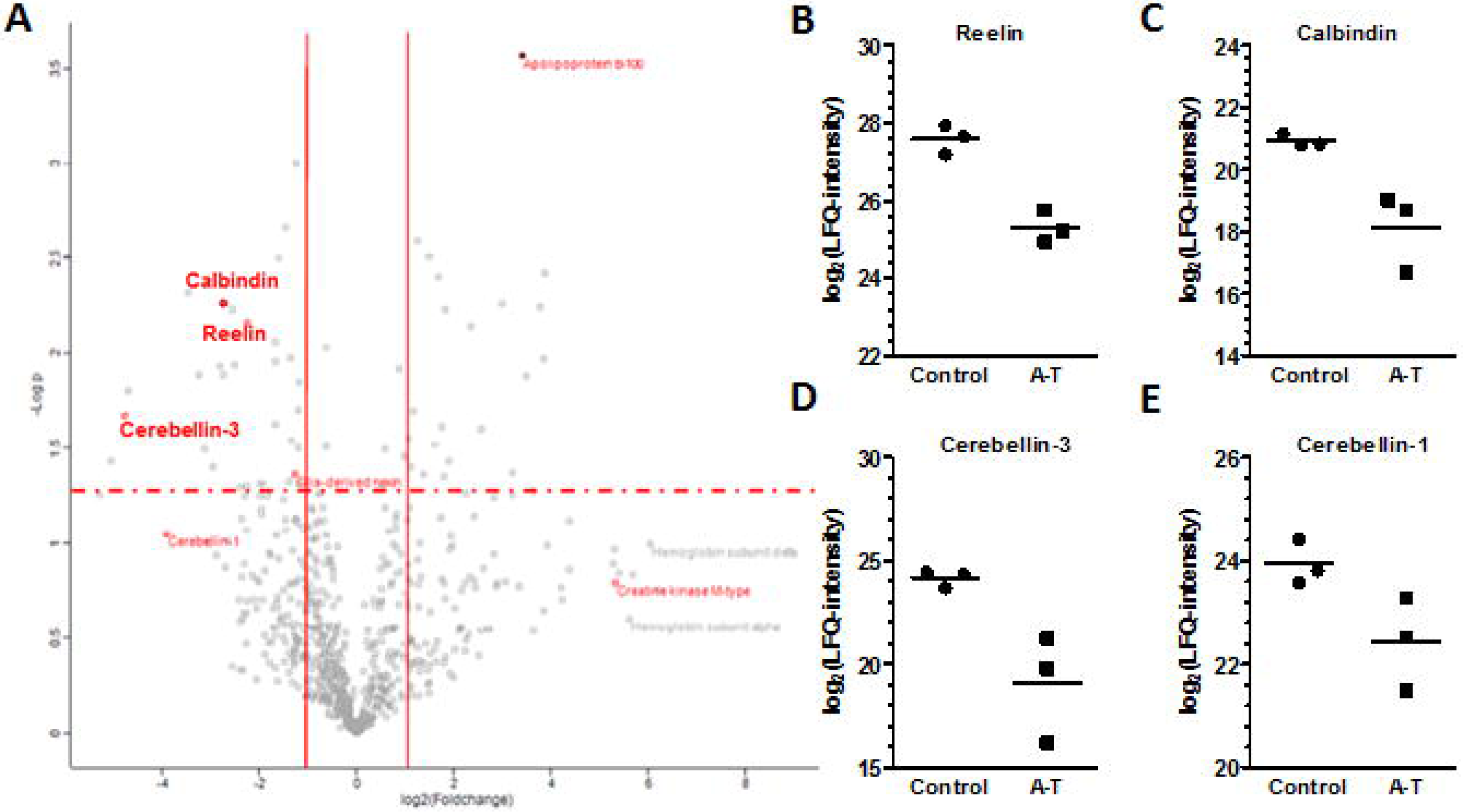
Proteomic analysis of the Cerebro-Spinal Fluid (CSF) collected from 3 oldest patients and controls (age 14-16 years). Analysis of the CSF of A-T samples was performed by a label-free protein quantification approach using nano ultra-high performance liquid chromatography/ nano electrospray mass spectrometry (nano UPLC/ nanoESI-MS). **(A)** Analysis of the data was performed with the MaxQuant quantitative proteomics software and illustrated by a Volcano plot. The red squares indicate proteins that display both large-magnitude fold-changes (x-axis) as well as high statistical significance (-log of p-value, y-axis). Label-free quantitation is shown for **(B)** Reelin, **(C)** Calbindin, **(D)** Cerebellin-3 and **(E)** Cerebellin-1 in A-T patients compared to controls.

As completely novel finding, a similarly strong downregulation was observed for Reelin (RELN, to 15%), which is expressed by the glutamatergic granule neurons of cerebellum [33]. The deficiency of Reelin could be upstream from PN degeneration in A-T, in view of previous knowledge: A spontaneous Reelin deletion in mouse (the famous “reeler″ mutant strain) causes ataxia due to brain cortex foliation defects from migration anomalies along radial glia, prominently affecting cerebellar PN [34,35]. In man, a rare autosomal recessive ataxia due to cerebellar hypoplasia with lissencephalopathy is caused by Reelin deficiency [36]. Reelin is a large secreted extracellular matrix glycoprotein that activates apolipoprotein receptors. The interaction between two APOE receptors (VLDLR and APOER2) and Reelin, as one of their ligands, controls not only neuronal positioning during brain development, but also synaptic plasticity in the adult brain [37,38]. APOE is a cholesterol transport protein and Reelin modulates intracellular cholesterol efflux in macrophages [39]. Reelin also influences postsynapses in their glutamatergic NMDA receptor composition, modulating postnatal neurogenesis, enhancing spine hypertrophy and long-term potentiation [40].

Reelin is very credible to be involved in ATM-triggered pathogenesis, for several reasons. Cerebellar Reelin is downregulated upon X-irradiation stress of embryos in the late gestation stage or upon corticosteroid exposure [41,42]. Reelin is expressed in glutamatergic granule neurons of the cerebellum, which secrete it from parallel fiber presynapses to PN dendrites and ensure spine development [33,43,44]. During adult life, Reelin is involved in the degeneration of PN [45]. Reelin signals are detected by the VLDLR and ApoER2 receptors, before they are intracellularly transduced in phosphorylation-dependent manner by DAB1 and the LIS1/PAFAH1B complex [46-48]. A mutation in PAFAH1B3 was reported to trigger ataxia and brain atrophy [49].

In excellent agreement with the observation of Reelin dysregulation, a significant downregulation was detected also for Fibroblast Growth Factor Receptor 1 (FGFR1, to 38%), which is expressed in the radial glia of cerebellum [50]. This trophic signaling molecule was already implicated in ATM-dependent genotoxicity response and in Reelin-dependent radial glia migration, with secondary effects on granule neurons and PN in the cerebellum [51-53]. The upregulation of Notch-3 (NOTCH3, to 260%) may be related to the same pathway, since NOTCH3 has been implicated in the role of Bergmann astroglia for the radial migration of neurons during cerebellar corticogenesis [54].

Further support for the central role of Reelin signaling in this proteome profile came from the massive upregulation observed for Apolipoprotein B-100 (APOB, to 1282%). This lipid transporter shows preferential expression in granule neurons and PN of the cerebellum according to the Allen mouse brain atlas (http://mouse.brain-map.org/gene/show/88416). The APOB/LDLR interaction activates the PI3K-PKCζ-Sp1-ABCA1 pathway in parallel manner to the Reelin/APOE-VLDLR/APOER2-DAB1 cascade in the control of cholesterol homeostasis [55]. The levels of APOB and the Reelin receptor VLDLR are known to correlate inversely [56]. Also the upregulation of Junction Plakoglobin (JUP, to 217%) is relevant here, since this adhesion factor interacts with DAB2 [57].

Finally, the significant upregulation of Apolipoprotein J (also called Clusterin, CLU, to 159%) in A-T patient CSF is relevant. Although the main isoform is a Golgi-localized molecular chaperone, a secreted isoform acts as ligand for the VLDLR and APOER2 receptor to signal via the Reelin pathway [58-60]. Furthermore, Apolipoprotein J is a known marker of PN pathology [61,62], its genetic variants play a role for hereditary dementia similar to variants in NOTCH3 [63], and its plasma concentration is associated with brain atrophy [64]. Moreover, the upregulations of APOB and CLU were accompanied by increased levels of Apolipoprotein H (APOH, to 157%) as a mediator of inflammatory changes [65], and of Serum Amyloid P Component (APCS, to 248%) as a suppressor of immune responses to abnormal DNA, known to genetically interact with APOE [66,67]. A relevant upregulation was also detected for Paraoxonase 1 (PON1, to 228%), which serves as universal factor of antioxidant defense [68]. PON1 detoxifies oxidized low density lipoprotein (LDL) [69], is secreted from cells with VLDL [70], is inhibited in activity by VLDL-associated triglycerides [71] and is associated with familial combined hyperlipidemia [72].

Thus, there is a clear enrichment of Reelin signaling and apolipoprotein homeostasis factors among the A-T CSF dysregulations.

### Protein interaction bioinformatics suggest glutamatergic axon input to PN as site of pathology, while identifying molecular biomarkers of neuroinflammation

In order to complement this literature-driven prioritization by additional unbiased bioinformatics, all significantly dysregulated proteins in Table 2 were screened for enrichments of specific biological processes, molecular functions, KEGG pathways and protein domains at the STRING webserver of the European Molecular Biology Laboratory in Heidelberg (Fig. 3). As assumed, significant protein-protein-interaction enrichment was observed for both the downregulations (p=0.00948) and upregulations (p=1.11e-15). Among the cellular components, enrichments for the extracellular region were detected for 36 downregulated (false discovery rate q=4.86e-16) and 20 upregulated proteins (q=2.11e-09), as expected for a study in CSF.

**Figure 3.**
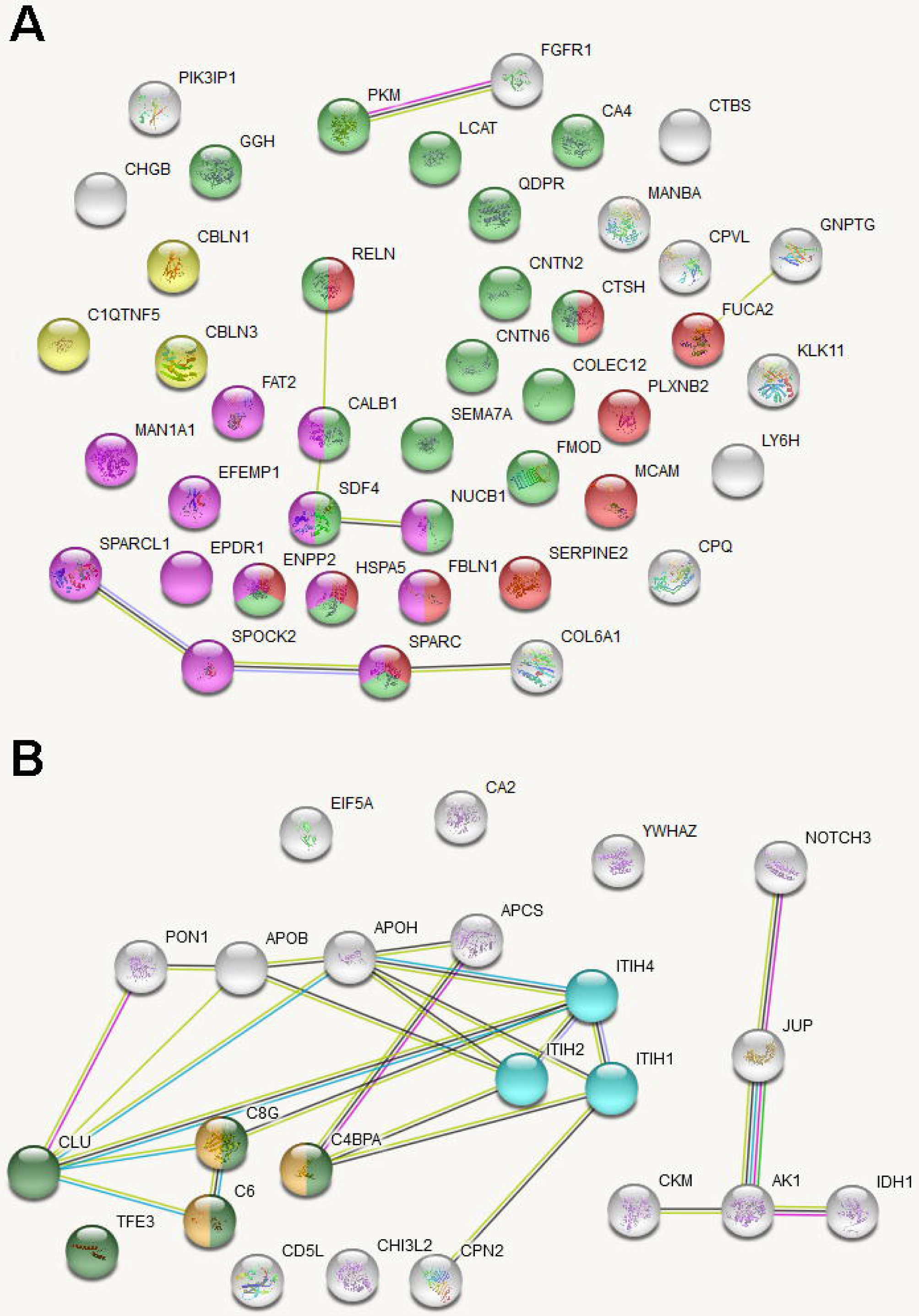
STRING diagram of protein-protein interactions and pathway enrichments in CSF of A-T patients. **(A)** The significant downregulations include pathway enrichments for “Calcium Ion Binding″ (magenta bullets), “Regulation of Locomotion″ (red bullets), “Response To Chemical″ (green bullets), “C1q domain″ (yellow bullets). **(B)** The significant upregulations include pathway enrichments for “Humoral Immune Response″ (forest green bullets), “Complement and Coagulation Cascades″ (gold bullets), “Hyaluronan Metabolic Process″ (cyan bullets). Previous knowledge of interaction between these factors from experiments, co-expression or text-mining is illustrated by connecting lines of various colors.

Among the 44 downregulations (Fig. 3A), novel specific insights included a significant enrichment for 18 factors in the “response to chemical″ (q=0.04, illustrated as light green bullets), among which RELN and SPARC (Secreted Protein Acidic and Cysteine Rich, to 49%) are involved in the response to irradiation [73,41], while CNTN2 (Contactin-2, to 78%) is involved in granule neuron radial migration [74,75]. A significant enrichment was observed also for 10 factors in the “regulation of locomotion″ (q=0.00962, illustrated as red bullets in Fig. 4A), among which SERPINE2 (Serpin Family-E Member-2, to 34%) regulates the differentiation of cerebellar granule neurons, while PLXNB2 (Plexin-B2, to 62%) is involved in their migration [76,77]. Significant enrichment was found as well for 13 factors in “calcium ion binding″ (q=2.38e-06, illustrated as purple bullets), among which the massive decrease of the protocadherin adhesion factor FAT2 (FAT Atypical Cadherin-2, to 3%) deserves special interest, since FAT2 is localized at parallel fibers of cerebellar granule neurons, is driving cellular motility, and its mutation causes Spinocerebellar ataxia type 45 [78-80]. Finally, a significant enrichment was observed for 3 factors with a “C1q domain″ (q=0.009, illustrated as yellow bullets), among which the downregulations of CBLN1 (Cerebellin-1, 19%) and CBLN3 (Cerebellin-3, 4%) were particularly strong. Both are secreted from parallel fibers of granule neurons and colocalize as heterodimeric complex with the glutamatergic GluRdelta2 receptor [81]. CBLN1 deficiency in mice was reported to cause cerebellar ataxia [82]. Further downregulations of relevance were detected for the motility factor ENPP2 (Ectonucleotide Pyrophosphate Phosphodiesterase-2, to 48%) [83], the PN marker SEMA7A (Semaphorin-7A, to 39%) [84], and the PN-expressed adhesion factor CNTN6 (Contactin-6, to 15%) that acts as NOTCH1 ligand [85,86]. CNTN6 deficiency in mice triggers ataxia [87]. In summary, markers for the glutamatergic parallel fiber afferents to PN, which originate in cerebellar granule neurons (such as FAT2 to 3%, CBLN3 to 4%, RELN to 15%, CBLN1 to 19%), appeared to be at least as diminished as PN markers (CALB1 to 15%, CNTN6 to 15%, SEMA7A to 39%). Overall, key factors of PN connectivity with their glutamatergic input axons are prominent among the downregulations.

**Figure 4:**
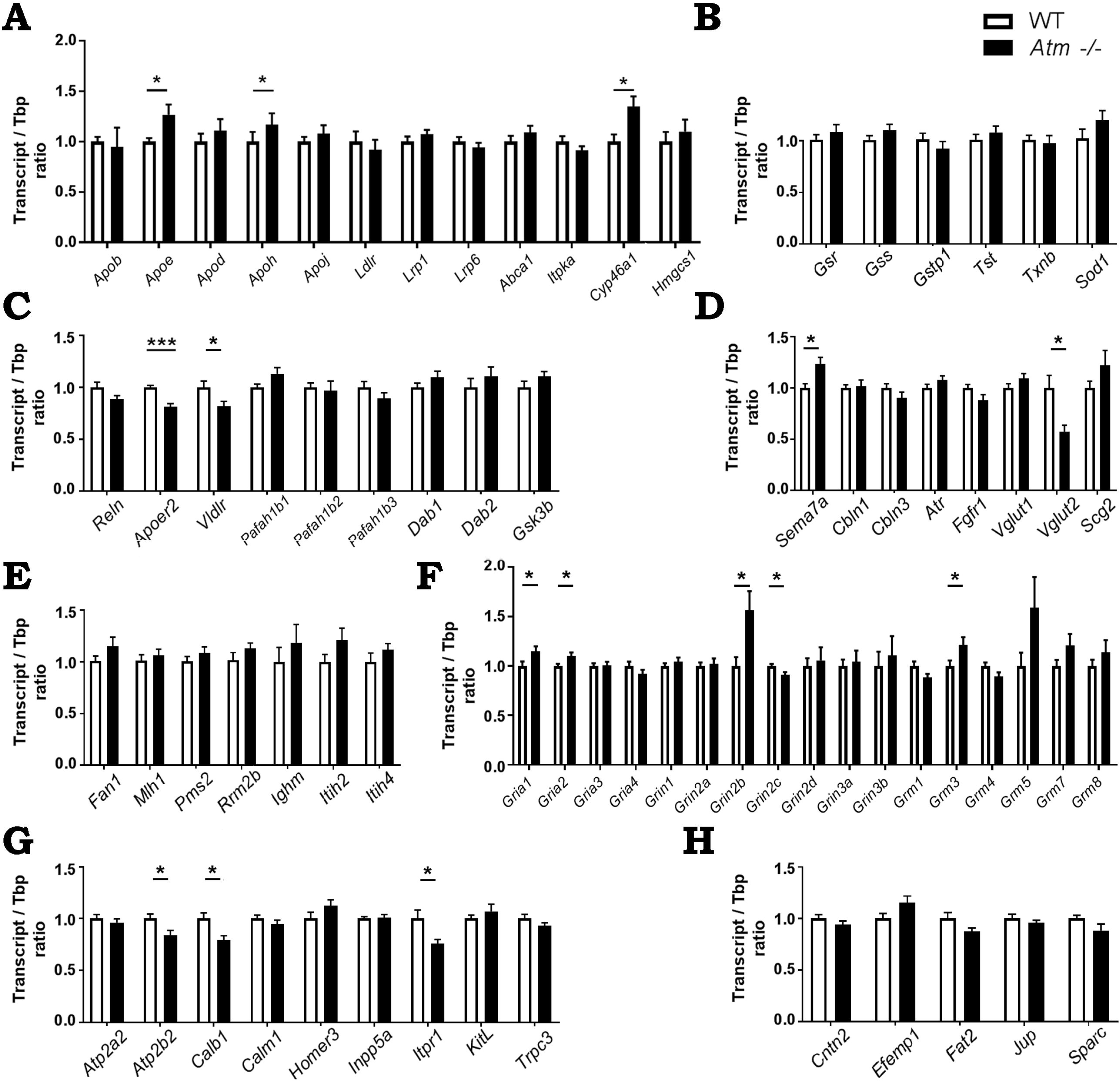
In cerebellar tissue from 2-month-old *Atm*-/- mice (black bars) versus Wild-Type mice (WT, white bars), the mRNA levels were assessed by quantitative real-time reverse-transcriptase PCR for crucial factors in several relevant pathways. **(A)** Among key factors of apolipoprotein and cholesterol trafficking, significant upregulations were observed for *Apoe, Apoh* and *Cyp46a1*. **(B)** No significant changes were detectable at this stage for any factors in the oxidative stress response. **(C)** Within the Reelin signaling pathway, the abundance of Reelin mRNA (*Reln*) was normal, but both Reelin receptors *Apoer2* and *Vldlr* showed significantly reduced transcript levels. **(D)** Among crucial modulators of the interaction between PN and afferent excitatory presynapses, an elevation was detected for *Sema7a*, whose gene product is secreted from PN to eliminate climbing fibers. In addition, *Vglut2* as a marker of climbing fibers showed a severe reduction. **(E)** Again, no dysregulation was documented at this stage for markers of DNA damage repair or neuroinflammation. **(F)** Glutamate receptor exhibited the expected elevation of the NMDA subunit *Grin2b* levels, which can be triggered by Reelin deficiency, as well as a decrease of *Grin2c* and increases for the AMPA subunits *Gria1* and *Gria2*, as well as the metabotropic subunit *Grm3*. **(G)** Among essential factors of calcium-dependent excitability, the significant downregulations of *Atp2b2, Calb1* and *Itpr1* clearly reflect incipient dysfunction of Purkinje neurons. **(H)** Despite various changes in PN postsyapses and glutamatergic presynapses, no alterations of the specific cell adhesion factors between cerebellar circuits could be found at this stage.

Among the 23 upregulations (Fig. 3B), novel specific insights included enrichments of 5 factors in the “humoral immune response″ (q=0.00007, illustrated as deep green bullets), among which CLU was already mentioned, while IGHM (Immuno-Globulin Heavy Constant Mu chain, to 2216%) is involved in acute inflammatory stress responses. Although not recognized by the automated bioinformatics, the upregulated factor CHI3L2 (Chitinase-3 Like-2, to 384%) is also involved in inflammatory responses [88,89], similar to the upregulated factors CPN2 (Carboxypeptidase-N Subunit-2, to 817%) [90] and the cholesterol biosynthesis suppressor CD5L (CD 5 Molecule Like, to 773%) [91-94]. Significant enrichment was found as well for 3 factors in the innate immunity “complement and coagulation cascades″ (q=0.017, illustrated as ochre bullets), among which C8G (Complement Component-8 Gamma Chain, to 195%) is part of the secreted lipocalin gene family and was implicated in the interleukin-6-mediated induction of acute-phase response [95,96], while C4BPA (Complement Component-4 Binding Protein A, to 143%) binds DNA to inhibit DNA-mediated complement activation and inflammatory responses against necrotic cells [97]. Significant enrichment was found also for 3 factors in the “hyaluronan metabolic process″ (q=0.008, illustrated as light blue bullets), namely ITIH1 (Inter-Alpha-Trypsin Inhibitor Heavy Chain-1, to 271%), ITIH2 (Inter-Alpha-Trypsin Inhibitor Heavy Chain-2, to 236%) and ITIH4 (Inter-Alpha-Trypsin Inhibitor Heavy Chain-4, to 213%). While the brain role of this gene family has not been thoroughly investigated, ITIH4 is induced by interleukin-6 during the acute phase response to infection [98,99]. Overall, neuroinflammatory responses are prominent among the upregulations.

### Validation in *Atm*-/- cerebellar mRNA confirms early Reelin signaling deficit

To assess whether these pathways have a prominent role also in cerebellar tissue of the A-T mouse model, we employed *Atm*-/- animals that were previously shown to die at ages of 4-6 months due to thymic lymphomas without showing signs of cerebellar degeneration. A hallmark of these mice is the high level of oxidative stress by Reactive Oxygen Species (ROS) that affects several tissues at the end of their lifespan, including the cerebellum [15]. Instead of quantifying secreted factors or studying their extracellular turnover, we analyzed transcript levels in 2-month-old cerebellar tissue for key components in several pathways, which were suggested by the A-T patient CSF proteomic profiles, namely cholesterol-apolipoprotein trafficking, Reelin signaling, glutamatergic input to PN, glutamate receptors, calcium homeostasis, oxidative stress, neuroinflammation and cell adhesion (Fig. 4).

As evidence for altered cholesterol trafficking and increased synthesis of ligands that may substitute for Reelin (Fig. 4A), significant increases were observed for the cholesterol-transporting apolipoproteins *Apoe* (to 127%, p=0.033), *Apoh* (to 159%, p=0.032) and the cholesterol efflux regulator *Cyp46a1* (to 135%, p=0.020), while markers for oxidative stress showed no dysregulation at this stage (Fig. 4B). As a crucial confirmation of the Reelin reduction in patient CSF, already these young *Atm*-/- cerebella showed significant mRNA decreases for both receptors of Reelin, namely *Vldlr* (to 82%, p=0.037) and *Apoer2* (to 82%, p=0.0003) upon qPCR (Fig. 4C). The transcriptional synthesis of the Reelin transcript (*Reln*) itself and its intracellular transducers *Pafah1b1, Pafah1b2, Pafah1b3, Dab1, Dab2* and *Gsk3b* appeared normal (Fig. 4C).

Analyzing key factors for the interaction of PN with excitatory afferents (Fig. 4D) demonstrated increased levels of the *Sema7a* (to 123%, p=0.009), a factor that is secreted from PN and contributes to the selective elimination of climbing fibers [84]. Indeed, a strong reduction was documented for *Vglut2* (to 57%, p=0.013), which serves as marker for climbing fibers and reflects the earliest pathology also in Spinocerebellar Ataxia type 1 [100]. There were no dysregulations for the Cerebellins (*Cbln1/Cbln3*) or *Fgfr1* at this stage (Fig. 4D), nor for any marker of DNA-damage repair and neuroinflammation under analysis (Fig. 4E). Consistent with the concept of altered lipid signaling and glutamatergic afferents, the glutamate NMDA receptor subunit *Grin2b* mRNA showed a strong upregulation (to 156%, p=0.034) (Fig. 4F). Given that Reelin levels are known to modulate GRIN2B levels in a reverse correlation during postnatal life [40,101], this *Grin2b* elevation supports the notion that Reelin signals are deficient in *Atm*-/- mouse cerebella already at the age of 2 months. In contrast, *Grin2c* was downregulated (to 91%, p=0.018), while upregulations were observed for the AMPA receptor subunits *Gria1* (to 115%, p=0.049), *Gria2* (to 110%, p=0.044) and for the metabotropic receptor subunit *Grm3* (to 121%, p=0.045) (Fig. 4F).

Among the calcium homeostasis factors, the significant decreases of mRNAs encoding the PN marker Calbindin-1 (*Calb1* to 79%, p=0.011), the Plasma Membrane Calcium ATPase (*Atp2b2* to 85%, p=0.024) and the ER-associated Inositol-1,4,5-Trisphosphate-Receptor-1 (*Itpr1* to 78%, p=0.026) were reflecting the deficient excitability of PN, downstream from Reelin and Glutamate signals (Fig. 4G). In comparison, none of the cell adhesion markers that were assessed showed any significant dysregulation (Fig. 4H). Thus, the transcriptional analyses confirm Reelin signaling deficits before the advent of altered locomotor behavior and PN death. Our data also indicate that the pathways of lipid and glutamate signaling with downstream calcium-dependent excitability are affected before DNA damage repair, oxidative stress responses, neuroinflammation and cellular adhesion show detectable changes. A diagram (Fig. 5) illustrates the scenario within the 2-month-old *Atm*-/- cerebellar circuitry, with red letters highlighting upregulated factors, while blue letters denote downregulations.

**Figure 5.**
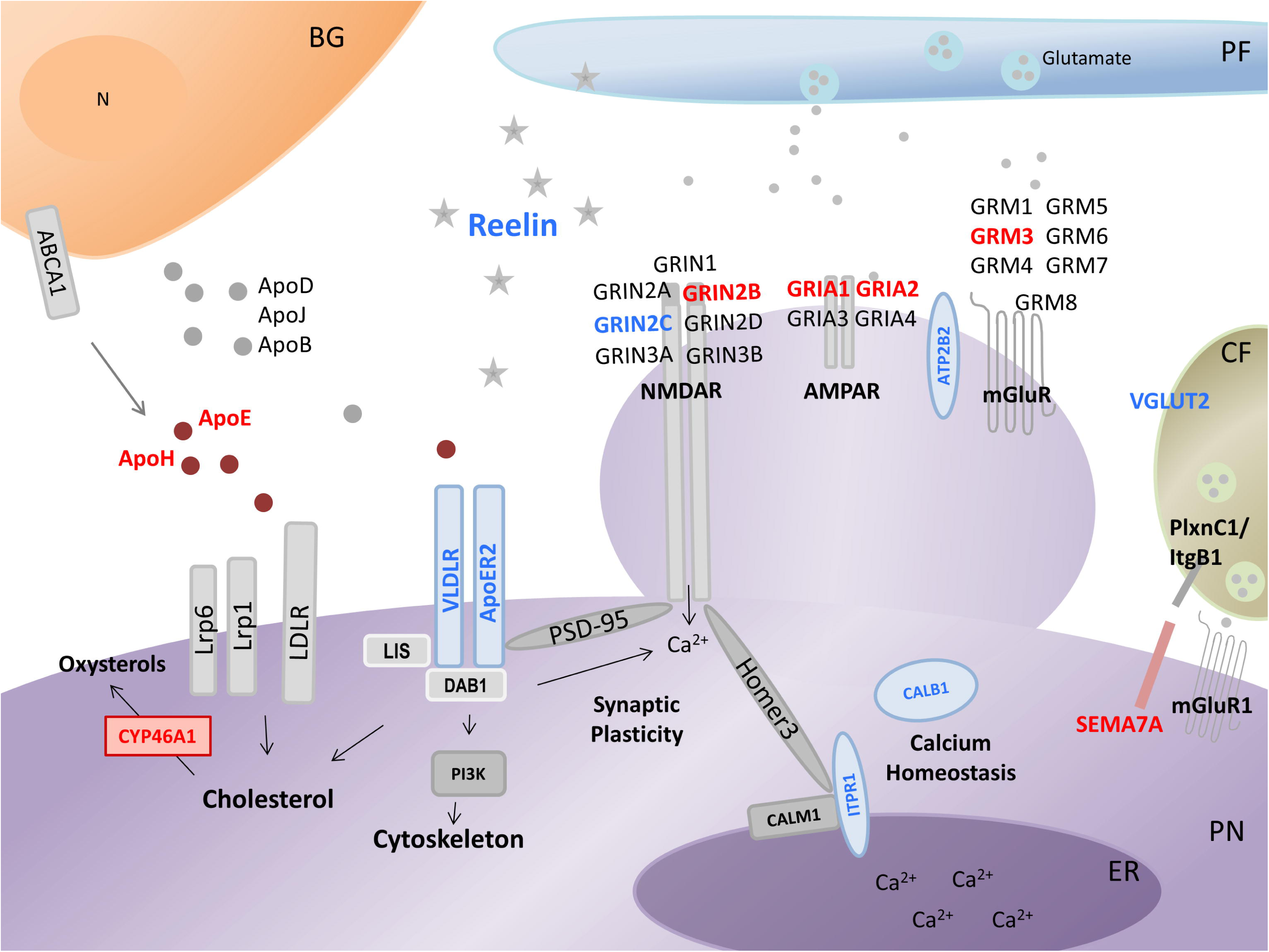
Schematic overview over apolipoprotein and glutamate signaling modulated via Reelin within neural circuits of the cerebellum (PN for Purkinje neuron, CF for Climbing Fiber, PF for Parallel Fiber, BG for Bergmann Glia, ER for Endoplasmic Reticulum, N for Nucleus). Relevant qPCR findings in the 2-month-old *Atm*-/- cerebellum were highlighted (red for upregulations, blue for downregulations).

## Conclusion

Comprehensive evaluations of pathogenesis pathways and of potential biomarkers have become possible via high-throughput sequencing technology. The advent of personalized medicine will make these approaches at DNA, RNA, protein and lipid level available soon in many hospitals to counsel families regarding disease risk, progression and therapy benefit. For A-T diagnostics, the levels of alpha-fetoprotein in blood extracellular fluid serve as well-established critical biomarker, so now for the first time the brain extracellular fluid was evaluated in a quantitative global screen. The CSF findings identified prominent changes of Reelin/apolipoprotein signaling and possibly adhesion at sites of glutamatergic input to PN, accompanied by neuroinflammatory activation, with surprising consistency. Our study therefore provides proof-of-principle that this approach is useful and should be pursued in further patients and genetic model studies.

The molecular dysregulations observed may be part of the irradiation and DNA damage repair pathways, given that Reelin and SPARC are modulated during radioresistance-reversal, while APCS and C4BPA interact with abnormal DNA. They may be part of neural migration and synaptic mobility, which certainly will involve extracellular factors that modulate adhesion, signal responses and locomotion. But importantly, the pathogenesis may also relate to altered glutamatergic excitation of PN dendrites. The ATM protein was recently shown to be a crucial facilitator of synaptic excitatory vesicle release, in a fine balance with the ATR protein that is responsible for inhibitory neurotransmission [102]. ATM levels rise in response to a blockade of NMDA receptors [102], so an influence on NMDA receptor composition and synaptic plasticity by Reelin signals [40,103] may play a key role in ATM deficiency. Dysfunction of the glutamatergic parallel fiber input is known to result in PN death [104]. Indeed, our transcriptional analyses in 2-month-old *Atm*-/- cerebella support the concept that Reelin signaling deficits are a novel, early and important feature in the pathogenesis of A-T neurodegeneration. Furthermore, our data suggest that the changes in signaling and excitability occur before responses to DNA damage, oxidative stress, neuroinflammation or cell adhesion deficits become notable.

To establish new approaches of neuroprotective value, it is important to note that the addition of recombinant Reelin to organotypic slice cultures via conditioned medium was observed to rescue the pathology of synaptic vesicles and of paired-pulse facilitation in reeler mutant mice [105]. In addition, the expression of Reelin and VLDLR can be upregulated by administration of the psychotropic drug olanzapine [106]. Thus, our observations may hold therapeutic value.

## Acknowledgements

This research received funding from the A-T Children’s Project. We thank Dr. Suzana Gispert-Sánchez and Katrin Krug for advice and technical assistance. We are also grateful to Konrad Bochennek for the collection of the CSF samples and give many thanks to Ramona Mayer for sample preparation for mass spectrometry.

